# Data quality of Whole Genome Bisulfite Sequencing on Illumina platforms

**DOI:** 10.1101/188797

**Authors:** Amanda Raine, Ulrika Liljedahl, Jessica Nordlund

## Abstract

The powerful HiSeq X sequencers with their patterned flowcell technology and fast turnaround times are instrumental for many large-scale genomic and epigenomic studies. However, assessment of DNA methylation by sodium bisulfite treatment results in sequencing libraries of low diversity, which may impact data quality and yield. In this report we assess the quality of WGBS data generated on the HiSeq X system in comparison with data generated on the HiSeq 2500 system and the newly released NovaSeq system. We report a systematic issue with low basecall quality scores assigned to guanines in the second read of WGBS when using certain Real Time Analysis (RTA) software versions on the HiSeq X sequencer, reminiscent of an issue that was previously reported with certain HiSeq 2500 software versions. However, with the HD.3.4.0/RTA 2.7.7 software upgrade for the HiSeq X system, we observed an overall improved quality and yield of the WGBS data generated, which in turn empowers cost-effective and high quality DNA methylation studies.

## Introduction

Methylation of cytosine residues (5-mC) in the CpG context is a key epigenetic mark which is involved in processes such as regulation of gene expression, cell differentiation, genomic imprinting, X-chromosome inactivation, transposon silencing and chromosomal stability [1]. Aberrant methylation patterns have been shown to be associated with a growing number of conditions and disease, in particular cancer [2]. Bisulfite conversion, in which unmethylated cytosines are converted to uracil (and subsequently to thymine after PCR) whilst methylated cytosines remain unchanged [3], remains the gold standard technique for detecting DNA methylation and is often used in combination with next generation sequencing for interrogation of methylation on a genome-wide scale. Whole genome bisulfite sequencing (WGBS) is the only method that provides an affordable and unbiased view of the entire methylome, which comprises ∼28 million CpG sites in humans.

Bisulfite converted libraries constitutes a particular case of low diversity sequences where the base composition is reduced to virtually three nucleotides (A,T,G) and a very small fraction of Cs, which represents the small portion of methylated cytosines in a genome. Sequencing of libraries with unbalanced base composition on the Illumina systems has historically been challenging, frequently leading to low data yields and inferior sequencing quality [4][5] [6]. Such issues can, in part, be attributed to earlier versions of the software operating the image analysis and base calling onboard the instrument, which were not adapted for handling low diversity sequences. The HiSeq Control Software (HCS) operates the imaging and calls the Real Time Analysis software (RTA) to execute intensity extraction, base calling and quality scoring. Previously, a dedicated control lane with a balanced library with A/T and G/C equally represented and a high level of spike-in with a high diversity library such as PhiX was required to improve data quality and yield [6]. For WGBS specifically, low base calling quality scores (Q-scores) assigned to guanines (representing methylated positions) have been reported in data generated with a certain version of HCS and RTA on the HiSeq 2500 system [7]. Moreover, the aforementioned WGBS data was shown to significantly deviate in terms of global methylation levels compared to data generated from identical libraries using other HiSeq 2500 software versions on the same instrument, suggesting that the reduced Q-scores of nucleotides representing 5-mC in bisulfite sequencing may result in biased methylation levels. Updates of the HCS and RTA software, which were first implemented on the MiSeq system and later also on the HiSeq2500 system (from HCS v2.2.38/ RTA1.18.61 and forward), conferred significant improvements in data yield and sequence quality of low diversity libraries, including bisulfite converted libraries [6].

Illumina’s HiSeq X system is the current work horse for whole genome sequencing studies. The patterned flow cell technology on the HiSeq X has significantly increased throughput and pushed costs close to the $1000 genome. The HiSeq X platform is now open to WGBS at an equal favorable price per base, and thus holds great promise for large-scale methylome studies. However, considering previous software related issues with bisulfite sequencing, an examination of the quality of WGBS data generated on the HiSeq X system is warranted. We therefore performed a retrospective quality control analysis of WGBS data that was generated from a set of control DNA samples within our core facility over time using different software versions on the HiSeq X, the HiSeq2500 and most recently, the NovaSeq. We identified substantial low Q-scores assigned to guanines in the second read of paired-end WGBS data with certain HCS/RTA versions. Notably, this issue was mitigated in the most recent HiSeq X software update HCS: HD.3.4.0 /RTA 2.7.7 and we demonstrate that the latest HCS/RTA version provide sequences of high quality, comparable to WGBS data generated on the HiSeq 2500 system. Despite low Q-scores assigned to guanines by certain software versions, we observed only minor differences in global methylation levels across libraries prepared with the same method. Rather, we observe that global methylation rates vary more depending on the choice of library preparation protocol. Albeit differences in global methylation rates, correlation of methylation at individual CpG sites across methods and sequencing software were in general high.

## Results and Discussion

Low Q-scores assigned to guanines in bisulfite sequencing reads generated with HiSeq 2500 systems installed with certain HCS/RTA software versions have previously been observed [7]. This particular issue was primarily observed at guanine positions in the second read (R2) of directional MethylC-Seq libraries whereas the first read (R1) did not significantly suffer from low Q-scores at any nucleotide type. The small fraction of cytosines, which are not converted during bisulfite treatment, represents methylated positions in the genome. Most types of WGBS library protocols are designed to sequence the original top and bottom DNA strands and thus methylated positions are sequenced as cytosines in R1 and as guanines in R2 of paired-end sequencing. Interestingly, a recent study demonstrated low Q-scores at guanines in R1 for PBAT-type bisulfite sequencing libraries, in a HCS/RTA software version specific manner [8]. In contrast to the majority of WGBS library protocols, PBAT libraries are constructed to sequence the complementary to the original strands, hence methylated cytosines are sequenced as guanines in R1 and as cytosines in R2 [8]. Low Q-scores assigned to guanines in R2 of MethylC-seq libraries or in R1 of PBAT libraries implies that the issue is related to low content of guanines in bisulfite converted reads. Moreover, as the fraction of guanines in R2 of most WGBS libraries (or R1 of PBAT) is very low, extensive Q-score reduction at guanines will not be obvious when inspecting average base quality in general. Overall low Q-scores of guanines in WGBS data may indicate a higher probability of base calling errors, which in turn could cause technical variance of methylation levels. Moreover, reduced Q-scores could potentially lead to lower alignment rates and data loss as less data might pass quality filtering. For bisulfite sequencing on HiSeq 2500 (v4 chemistry), the guanine Q-score issue was mitigated from software update HCS v2.2.38/RTA 1.18.61 and forward so that WGBS data generated with this system is now generally of high quality [9] and can such be used for comparison with data generated on the HiSeq X and NovaSeq systems.

In order to retrospectively examine the quality of bisulfite data generated with the HiSeq X system we analyzed a set of WGBS libraries from a core set of DNA samples that were prepared with three different library preparation protocols and sequenced across different software versions. The WGBS data was generated with three different HCS and RTA versions for HiSeq X; HCS v3.3.39/RTA 2.7.1, v3.3.75 /RTA 2.7.5, and the most recent update; HD.3.4.0 /RTA 2.7.7. In addition we included WGBS data generated from a pilot sequencing run on the NovaSeq instrument. For this comparison we examined sequence data that were generated from Accel-NGS Methyl Seq (Swift Biosciences), TruSeq DNA Methylation (TSDM, formerly EpiGnome, Illumina Inc) and <u>spl</u>inted ligation <u>a</u>dapter <u>tagging</u> (SPLAT,[10]) libraries prepared from lymphoblastoid cell lines (NA10860, NA11992) or the leukemic cell line REH. The DNA samples used, library pools and sequence data generated is outlined in S1 Fig. General metrics for all of the sequencing runs are found in Table 1. For comparison we included previously published data generated from libraries from the same DNA samples sequenced on the HiSeq 2500 system (HCS v2.2.38/RTA 1.18.61) [9].

## Low quality scores assigned to guanines in R2 by certain HiSeq X HCS/RTA software versions

To assess the quality of data generated by different HCS/RTA versions on HiSeq X we considered Q-scores assigned to the four nucleotide types separately (A, C, G and T). On average high Q-scores were assigned to all the four nucleotide types in R1 (average Q-score per nucleotide type: 33-38), irrespective of sample, library protocol, or software version on the sequencer (S1 and S2 Tables). With HCS v3.3.39/RTA 2.7.1 and v3.3.75/RTA 2.7.5 the Q-scores assigned to adenine, cytosine or thymine bases in R2 were generally lower than in R1, although still on average > 30. Notably, libraries sequenced on HiSeq X systems installed with the aforementioned software versions displayed significantly lower and more variable Q-scores at guanines in R2 compared to the other nucleotide types, even though high amounts of a balanced PhiX library was spiked-in to each lane (20-40%).a This feature was observed in all library types and samples, however it was particularly pronounced in data from SPLAT libraries (average G-base Q-scores: 22-29) in comparison to data generated with TSDM libraries (average: 26-32). (Figs 1 and 2, S1 Table). SPLAT libraries prepared from various human sample sources (REH, NA10860, NA11992), each with differing levels of global methylation, were equally affected (S1 Table). TSDM libraries display a typical and pronounced GC bias (average 26.1-26.8% GC content in bisulfite converted reads for NA10860), meaning that GC regions are overrepresented in this library type, whereas SPLAT libraries have a more even representation of the genome average (22.5-25.1%GC content in bisulfite converted reads for NA10860) [9]. Therefore, it appears that the guanine Q-scores assigned by RTA: 2.7.1 and RTA:2.7.5 appear to improve with increasing GC content of the library (S2 Fig).

**Figure 1.**
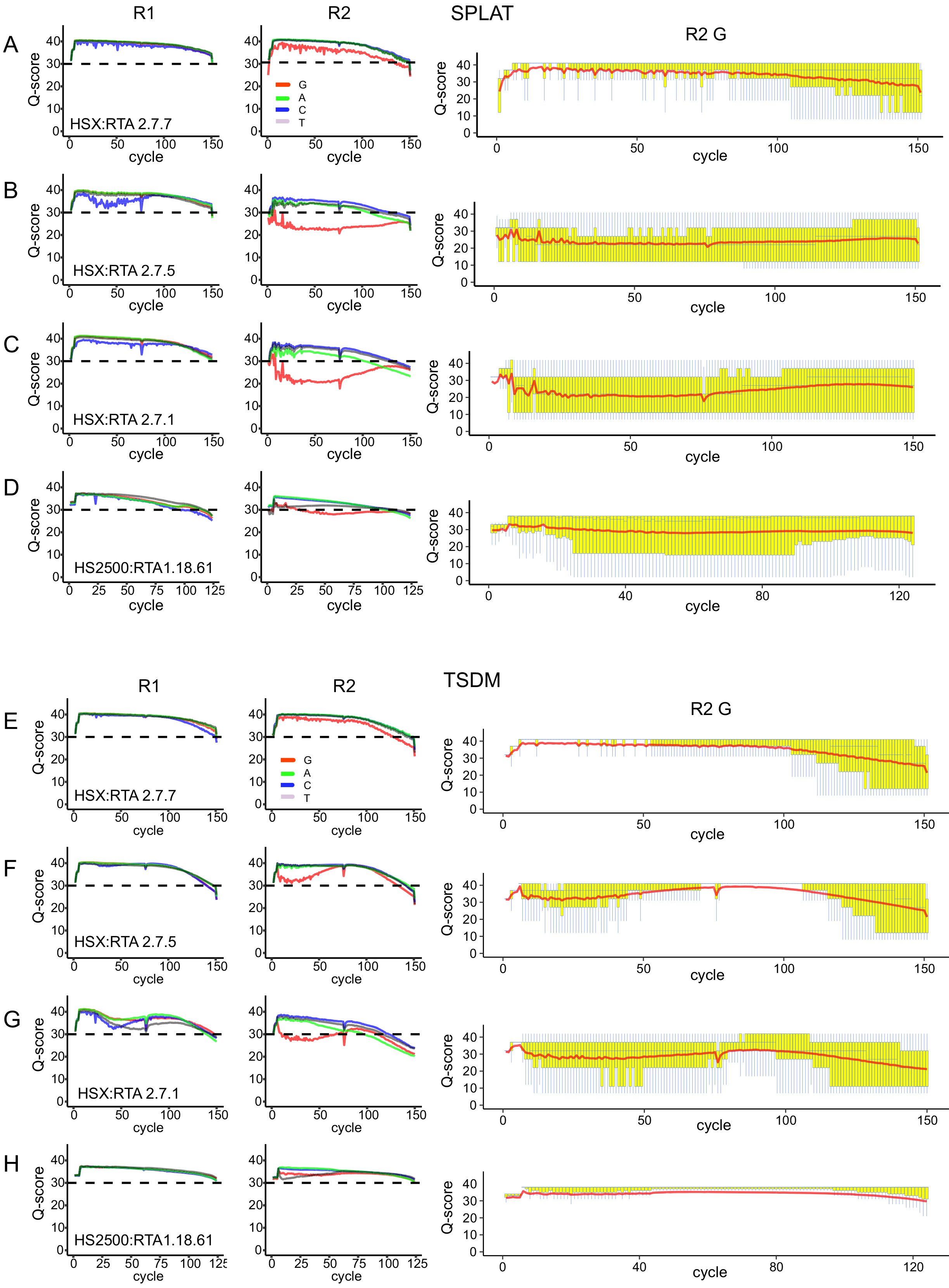
Examples of average base call quality scores for whole genome bisulfite sequencing of libraries prepared from lymphoblastoid cell line NA10860. Per nucleotide quality scores (average for each sequencing cycle) for read 1 and read 2 separately. A-D) SPLAT libraries. E-H) TSDM libraries. Panels A,B,C and E,F,G show Q-scores obtained with HiSeq X RTA versions, the version numbers are noted in each panel. Panels D and H show corresponding data generated on the HiSeq 2500 platform. Q-boxplots for guanines exclusively in read 2 are plotted in the rightmost panels.

**Figure 2.**
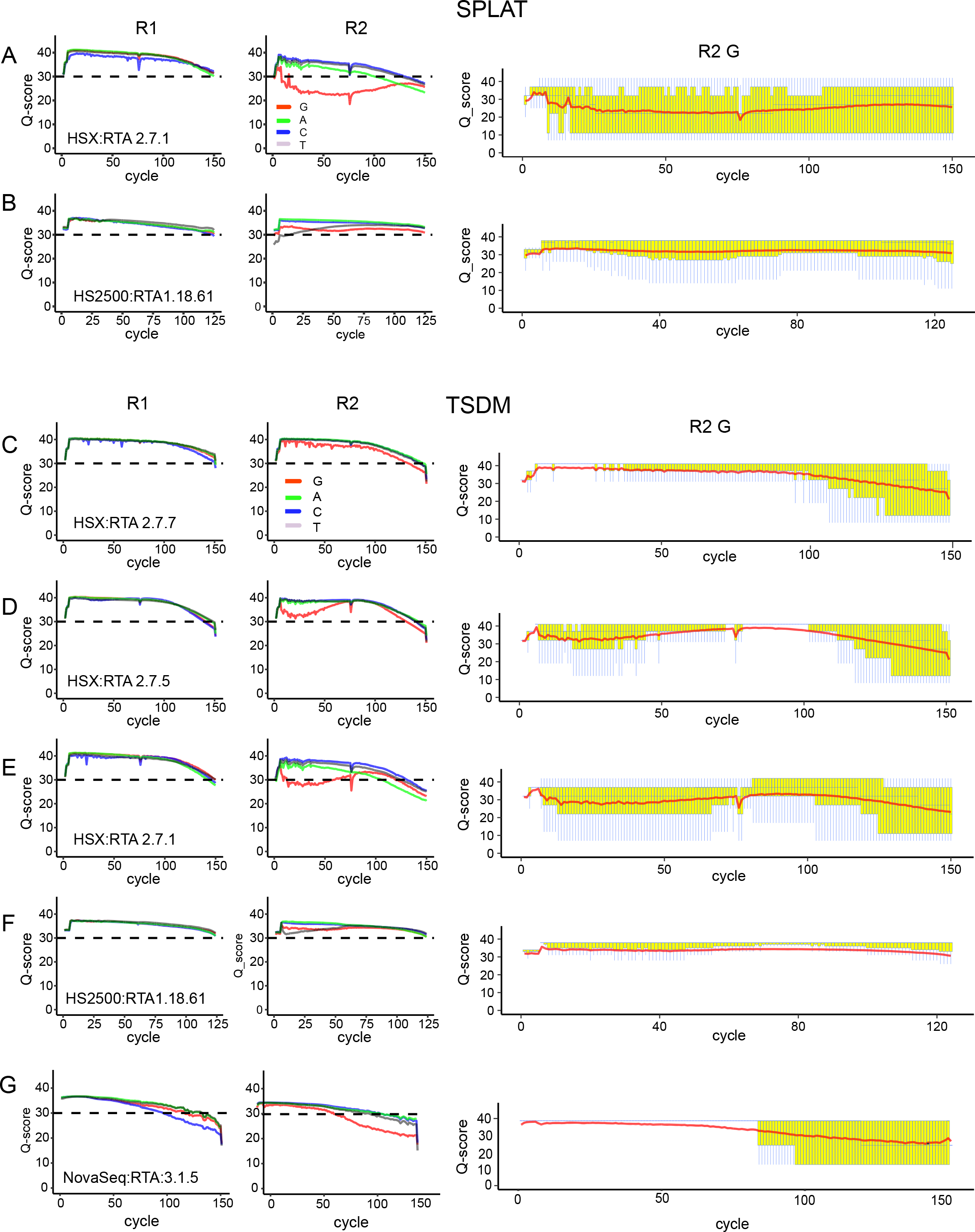
Examples of average base call quality scores for whole genome bisulfite sequencing of libraries prepared from leukemia cell line REH. Per nucleotide quality scores (average for each sequencing cycle) for read 1 and read 2 separately. A and B): SPLAT libraries. C-F) TSDM libraries. HiSeq X RTA versions are plotted in Panels A, and C-E. Corresponding data generated on the HiSeq 2500 are plotted in panels B and F and data generated on the NovaSeq are plotted in panel G. Q-boxplots for guanines exclusively in read 2 is plotted in the rightmost panels.

## WGBS data quality and yield with the HiSeqX HD.3.4.0 /RTA 2.7.7 software update

The HCS/RTA software update for the HiSeq X system (HD.3.4.0 /RTA 2.7.7) was released by Illumina in February 2017 in order to achieve better performance of WGBS on the HiSeq X. For this software version, WGBS data was available for three different WGBS library types (TSDM, SPLAT and Accel-NGS) to an average of ~500 M read pairs per library with a 2%PhiX spiked-in (S1 Fig).

Generally, all the libraries displayed high Q-scores for all four nucleotides types in both reads pairs, which were consistent across all sequencing lanes at similar levels to WGBS data generated from the same library types and DNA samples on the HiSeq 2500 (Figs 1 and 2, S3 Fig and S2 Table). Notably, the same exact SPLAT library that resulted in low Q-scores when sequenced with HCS v3.3.39/RTA 2.7.1(SPLAT-1a,average guanine Q-score:22) exhibited much improved quality when the same library was sequenced with software version HD.3.4.0 /RTA 2.7.7 (SPLAT-1a, average guanine Q-score: 32-33). This finding strengthens the notion that the Q-score issue is software related and does not originate from the sequencing library per se. Moreover, the trend towards higher guanine Q-scores with libraries with increased GC content, which was observed with the earlier HiSeq X software versions, was not as evident in the data generated by HCS/RTA software version HD.3.4.0 /RTA 2.7.7 (S2 Fig).

Next, we were interested in how the amount of “usable” data (post alignment) obtained from a HiSeq X lane compares to a HiSeq2500 lane. Alignment rates of the WGBS data generated on the HiSeq X with HD.3.4.0/RTA 2.7.7 were on par with previously generated WGBS data from HiSeq 2500 [9] and higher than those obtained from the previous HiSeq X software version (77-80% as compared to 65-75%) (Table 1 and S3 Table). The levels of duplicate reads were 15-20%for SPLAT and Accel-NGS Methyl-Seq,whilst the TSDM libraries had higher (34-40%) duplication rates. All of the libraries displayed higher duplication rates on the HiSeq X than what was observed for the same library types sequenced on HiSeq 2500 (~2% and 15% for SPLAT and TSDM respectively). A large part of the duplicated reads in the SPLAT and Accel-NGS libraries are likely ExAmp duplicated reads, although we did not observe any obvious difference in the level of duplicate reads when loading different amounts of library of on the flow cell (100, 150 or 200 pM) (Table 1). Following adapter and quality trimming, mapping and deduplication,57-61%of the generated raw data was retained for the SPLAT library, 50-52%for the Accel-NGS Methyl-Seq library and 34-40%for the TSDM libraries (Table 1).Thus, more data was retained for SPLAT and Accel-NGS Methyl-Seq libraries, as a result of longer insert sizes and lower duplication rates compared to the TSDM libraries. On average we obtained data (post mapping and deduplication) corresponding to ~25x genome coverage per HiSeq X lane for SPLAT, ~22x for Accel-NGS Methyl-Seq, and ~17x for TSDM. Thus from one HiSeq X lane we generated approximately the same amount of high quality WGBS data as was previously obtained from two HiSeq2500 lanes [9], at approximately one fourth of the cost.

**Table 1.**
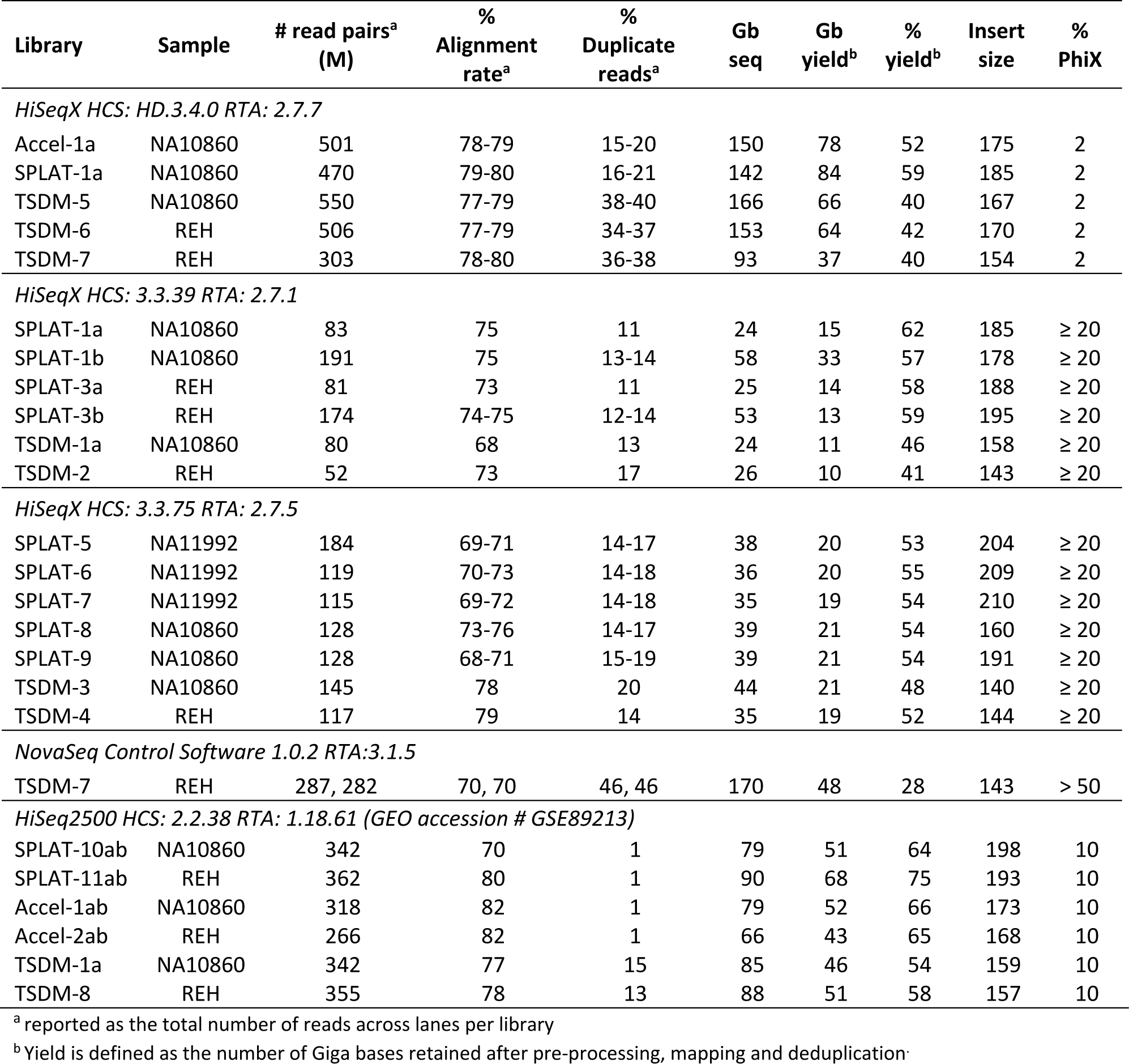
Per library sequencing metrics.

## WGBS on NovaSeq; results from a pilot experiment

NovaSeq is the latest iteration of Illumina sequencers for which the patterned flow cell technology is paired with a 2-color detection system for enhanced speed and throughput. For an initial assessment of WGBS data generated on the NovaSeq system we ran a single TSDM WGBS prepared from the REH cell line (TSDM-7), which was also sequenced on the HiSeqX with HD.3.4.0 RTA: 2.7.7. The library was sequenced on a single S2 flow cell across two lanes together with a pool of well-balanced libraries corresponding to 12% of the data derived from each lane. The remaining fraction contained PhiX (50%, installation run) and RNA-seq libraries. We observed a tendency towards lower Q-scores at the end of reads, which was most prominent for the methylated bases (C in R1 and G in R2, Fig 2G). The alignment rate was lower (70%) than obtained for the identical TSDM-7 library sequenced on HiSeqX (80%), most likely as a consequence of the lower Q-scores (Table 1). High duplication rates and lower mapping efficiencies resulted in a total yield of 28% of the unprocessed data compared to 40% for the identical TSDM-7 library sequenced on HiSeqX (RTA2.7.7).

## Minor variation in average methylation levels between data derived from different HCS/RTA software versions

In a previous study, up to 5%deviation in global methylation levels was observed in PBAT WGBS libraries sequenced with HCS 2.0.12/RTA 17.21.3 version on the HiSeq 2500, presumably due to low Q-scores of guanine bases that correspond to methylated cytosines [7]. Thus, in a similar fashion, we compared global methylation levels across data generated on the HiSeq X with the various software versions and HiSeq 2500 in order to determine if the low Q-scores is associated with bias in methylation-calling ((Fig 3A, S3 Table). Notably, within the same DNA sample and library type we observed only negligible (min-2%) differences in global methylation levels, which does not appear to be software-specific. For example, the largest difference observed was 1.6%between SPLAT-NA10860 libraries produced with the original protocol, sequenced on the HiSeqX RTA version HD.3.4.0 /RTA 2.7.7 in comparison to the HiSeq2500 (S3 Table). For SPLAT libraries amplified with a different PCR polymerase (Phusion U) the largest difference observed was 2%.As discussed previously, the lowest guanine Q-scores in R2 were observed in SPLAT libraries when sequenced on HCS v3.3.39 /RTA 2.7.1. Thus, we analyzed the global methylation levels of R1 and R2 independently in these libraries, and found that the largest difference observed was only a 0.5%difference in global methylation (Table 2).

**Table 2.** **Global methylation levels computed from R1 and R2 separately**

| Library | Sample | Software version | Average methylation R1 | Average methylation R2 | SR Alignment rate R1 | SR Alignment rate R2 | PE Alignment rate |
| --- | --- | --- | --- | --- | --- | --- | --- |
| SPLAT-1a | NA10860 | HCS: HD.3.4.0<br>RTA:2.7.7 | 55.5% | 55.8 % | 86.1 % | 83.7 % | 80.3 % |
| SPLAT-1a | NA10860 | HCS: 3.3.39<br>RTA:2.7.1 | 55.6 % | 56.1% | 82.5 % | 77.4 % | 75.3 |
| SPLAT-1b | NA10860 | HCS: 3.3.39<br>RTA:2.7.1 | 55.6% | 56.1 % | 82.6% | 76.6% | 74.8 |

On the other hand, the different library preparation protocols resulted in up to 4.5%differences in global methylation levels. The difference was most prominent in the immortalized B-cell line, NA10860, which is intermediary methylated on a global scale (average per library 55.8-60.3%). The highest global methylation levels derived from the NA10860 cell line were observed in the TSDM libraries (59.1-60.3%) and lowest in SPLAT and Accel-NGS Methyl-Seq libraries (55.8%-57.7%) (Fig 3A, S3 Table). Analysis of average methylation in 100 kB windows in the same data further confirmed this pattern (Fig 3B). For the leukemic REH cell-line, the largest inter-library difference observed was 2.6%(average global methylation levels 81.1-82.1% and 80.3-80.6%for SPLAT/Accel-NGS and TSDM libraries, respectively). Hence, we suggest that the variability observed in global methylation levels are more likely caused by alternative factors in the chemistry or PCR amplification used during library preparation than by issues with base calling or Q-scores and may be related to the GC biases observed for some library methods.

**Figure 3.**
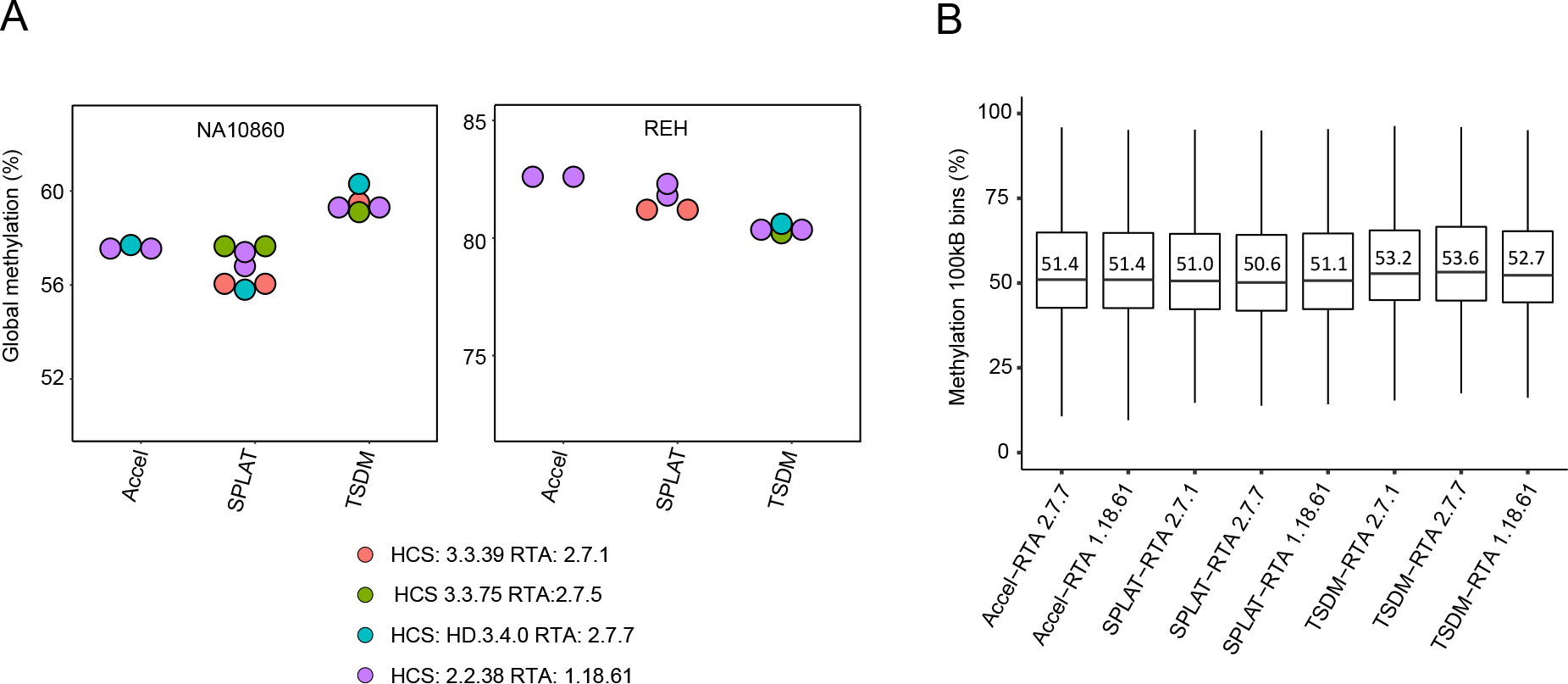
Variation in global methylation rates depends less on RTA version than on library preparation method. A) Global methylation levels (average methylation level across the whole genome) for DNA samples NA10860 and REH are shown for the different library preparation methods and are colored according to RTA software. B) Boxplots showing the average methylation in 100 kB windows for the various libraries and RTA versions; the median values are denoted in the panel.

## Concordance of methylation calls between HiSeq platforms

Next, we analyzed the correlation of methylation calls across the same type of libraries sequenced on HiSeq X and HiSeq 2500, by comparing either individual CpG sites or average methylation in 100 kb non overlapping windows. For methylation in 100 kb windows the Pearson’s correlation coefficient (Pearson’s R) was > 0.99 in comparisons between HiSeq X (HCS: HD.3.4.0 /RTA:2.7.7) and HiSeq 2500 verifying that methylation levels are called with high confidence on the HiSeq X platform ( Fig 4A). Similarly, the Pearson’s R was > 0.99 when we compared the average methylation in 100 kB windows between all versions of the HiSeqX and HiSeq2500 (data not shown).

**Figure 4.**
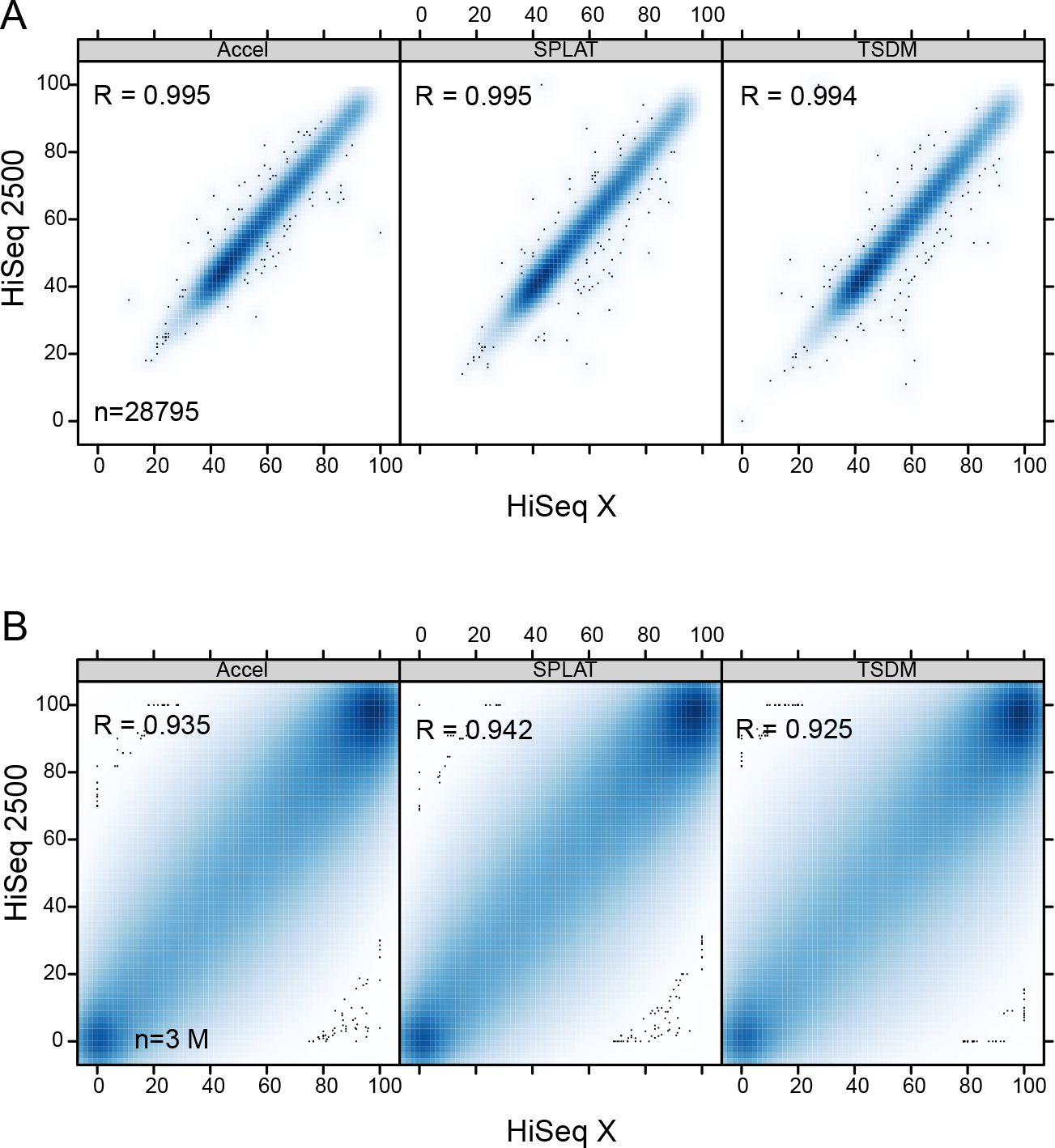
Scatter plots illustrating the high correlation of methylation calls. A) Average methylation in 100 kB windows (n=28,795) shown for data generated on HiSeq X and HiSeq 2500 systems. B) Methylation at individual CpG sites covered by more than 10 reads (n=7.5 M) shown for the corresponding data sets.

In order to obtain sufficient coverage to assess correlation at individual CpG sites across the various datasets we combined technical replicates sequenced with the same software versions (SPLAT1a+b, SPLAT3a+b, Accel1a+b, etc, see S1 Fig for an overview). In pairwise comparisons of single CpG sites, only sites covered by >10 reads, in each of the libraries were included. This resulted in the analysis of ~3 M CpG sites in total and thus the correlations were computed across the same set of CpG sites in all comparisons. The correlations varied depending on sample type (Table 3), however across high Q-score runs (HiSeq X RTA 2.7.7 vs HiSeq2500 RTA 1.18.61), the Pearsons’s R was 0.92‐0.94 for intra-library comparisons of the NA10860 sample and 0.96 for the REH sample (Fig 4B and Fig 5A). When comparing SPLAT libraries sequenced with software versions displaying low guanine Q-scores in R2 (SPLAT1ab for NA10860 and SPLAT3ab; RTA2.7.1) to those with high quality scores (SPLAT 1a; RTA2.7.7 and SPLAT10ab, SPLAT11ab; RTA 1.18.61) the Pearson’s R was 0.93 and 0.96(for NA10860 and REH respectively) and thus in the same range as the high Q-score runs Fig 5A). The highest correlation was observed for the TSDM-7 library that was sequenced on both HiSeqX (RTA2.7.7) and NovaSeq (Pearson’s R=0.98). For intra?library comparisons (e.g SPLAT-TSDM, SPLAT-Accel etc) the correlation coefficient was in the same range as the intra-library comparisons (sample NA10860; Pearson’s R=0.92-0.94), irrespective of software version (Fig 5A). As a second mean to measure the variability of methylation at individual CpG sites across data we computed the pairwise root mean square error (RMSE). The RSME values were generally very low within the same DNA sample type; 0.031-0.052 for the REH sample and 0.057-0.069 for the NA10860 sample (Fig 5B). Thus, we conclude that low guanine Q-scores in R2 does not have significant bearing on the CpG site methylation levels across the data sets analyzed herein.

**Figure 5.**
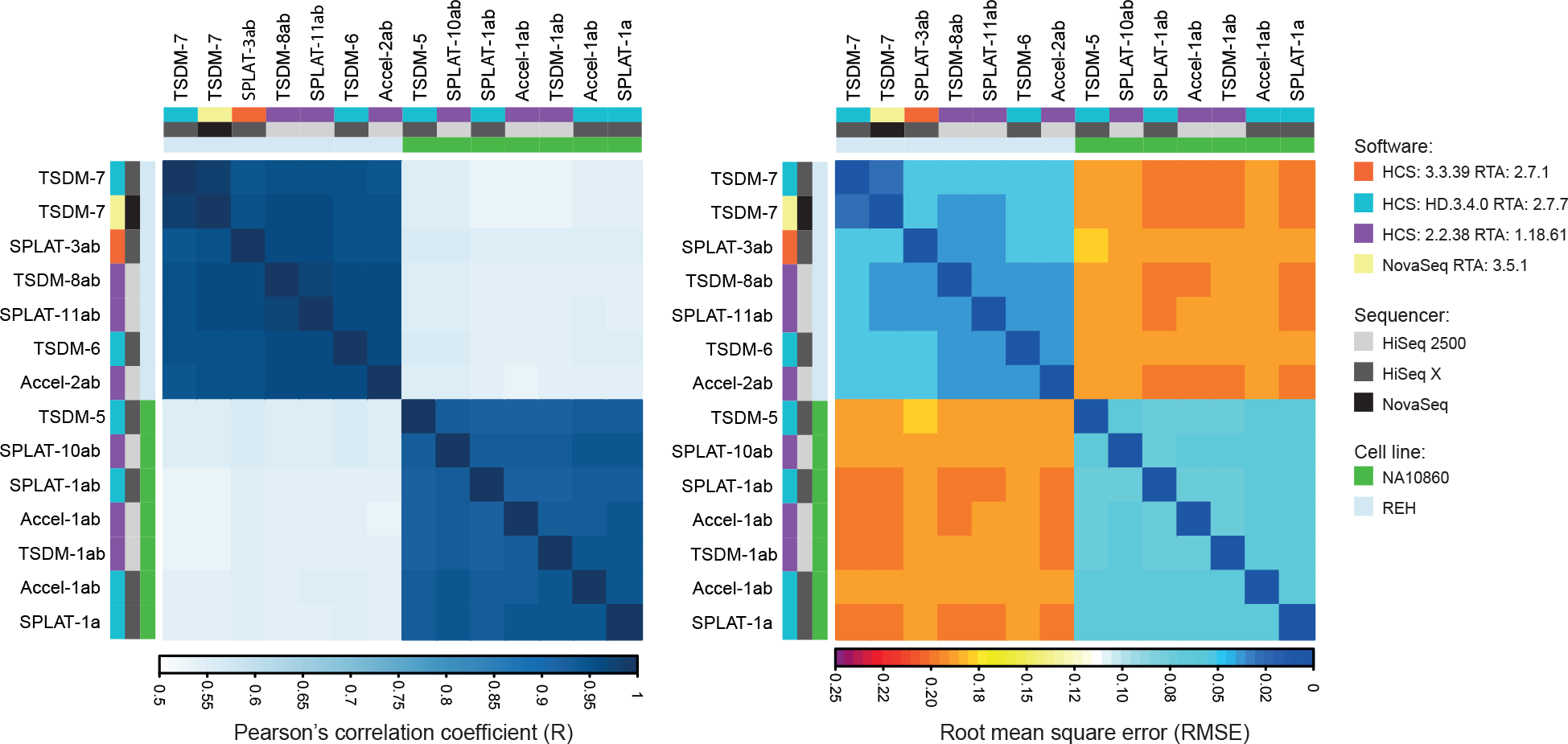
Correlation plots showing pairwise comparisons across a shared set of 3 M CpG sites covered by 10 reads or more in each dataset. A). Pearsons correlation coefficient for comparisons across all the library types, sequencing softwares, and cell types used in the study. B) The corresponding root mean square error (RMSE) values for the comparisons shown in panel A.

## Conclusion

In this report we compared WGBS data generated with different Illumina platforms and software versions. We found that certain software versions on the HiSeq X exhibit severe issues with base quality scoring exclusively at guanines in R2, which represent methylated positions. In such instances where the Q-tables used by the base calling software is not adapted to bisulfite reads it is not possible to determine if a low Q-score value reflects a true uncertainty in the guanine base calling or whether the base quality scores are merely under-estimated.

However, as we demonstrate in the present study, when using a popular workflow for WGBS pre-processing, alignment and methylation calling [11] we did not detect any gross methylation bias in those WGBS datasets with low R2 guanine Q-scores as compared to corresponding data with higher Q-scores. Although software-related methylation biases cannot be completely ruled out by our comparison, it is important to note that larger differences in absolute methylation levels were observed across library preparation methods than between different software versions. Nevertheless, we advocate that inspecting base quality scores per nucleotide type for WGBS generated on Illumina systems should be the standard and that sequencer software version should be reported for WGBS data submitted to journals and databases. Importantly, although many WGBS libraries still suffer from short insert sizes and high read duplication levels resulting in comprehensive data loss, with respect to base call quality we find that WGBS data generated with the HD.3.4.0 /RTA 2.7.7 HiSeq X version generated high quality data comparable to those obtained with HiSeq2500, at approximately one fourth of the per base cost.

We also evaluated WGBS data from an installation run on the new NovaSeq system and found that the present Q-scoring is still not optimal for bisulfite sequencing. Despite this, the methylation concordance was very high when comparing to data generated on the HiSeq X system. In our installation run, 50% phiX was spiked-in to evaluate the performance of the machine, but it should be noted that high amounts of phiX or other balanced library is not likely going to be required for WGBS on the NovaSeq. Additional data are needed to systematically assess and properly validate how the two-color detection system and the novel Q-score binning approach applied on NovaSeq (using only four Q-score values) reconcile with bisulfite sequencing and downstream analysis.

## Materials and Methods

### Library preparation

Human genomic DNA from lymphoblastoid B-cell lines was obtained from the Coriell Institute for Medical Research. Genomic DNA from the pre-B acute lymphoblastoid leukemia cell line REH (was isolated using the AllPrep Universal kit (Qiagen).

The EZ DNA Methylation Gold kit (Zymo Research) was used for sodium bisulfite conversion of DNA prior to library preparation. All WGBS libraries were prepared from 100 ng of genomic DNA. Accel-NGS Methyl-Seq (Swift BioSciences) and TruSeq DNA Methylation (Illumina Inc) were prepared according to the manufacturer’s protocols. SPLAT libraries were prepared as described previously [9] with the exception for libraries SPLAT-8 and SPLAT-9 which were amplified with the Phusion U Hot Start DNA polymerase (Thermo Fisher Scientific), whereas the other SPLAT libraries were amplified using the KAPA HiFi Uracil+ DNA polymerase (KAPA Biosystems).

### Sequencing and data analysis

Paired end sequencing (2 × 150) was performed on a HiSeq X system at the SNP&SEQ Technology Platform. The amount of library loaded on the instrument varied between 100 and 200 pM. A PhiX library was spiked in at 20-40% for HCS v3.3.39 /RTA:2.7.1 or v3.3.75/RTA:2.7.5 and at 2% for HD.3.4.0 /RTA:2.7.7). For comparison we also analyzed previously generated sequencing data from the HiSeq2500 system (HCS 2.2.38 / RTA 1.18.61) using the TruSeq v.4 chemistry PE125 (10 % PhiX) [9] and data generated on the an installation run of a NovaSeq 6000 instrument with 50% phiX spike-in (RTA:3.1.5).

Per nucleotide quality scores were extracted using Fastx v 0.0.14 or reported by Sisyphus, an in-house pipeline used at the SNP&SEQ Technology Platform for processing and QC of Illumina sequence data. Sequence reads were quality filtered and adaptors were trimmed using TrimGalore. For Accel-NGS Methyl Seq libraries 18 bp was trimmed off the 5’-end of R2 and the 3’ end of R1 to remove bases derived from the sequence tag introduced in the library preparation procedure. Alignment to the human reference assembly GRCh37 and methylation calling was performed with the Bismark software [11] and the pipeline tool ClusterFlow [12]. For TSDM libraries the initial 6 base pairs of each reads were ignored in the methylation calling procedure, to avoid random priming biases. Global methylation rates (ΣC/Σ(C+T) in CpG context) and methylation at individual CpG sites was obtained from Bismark methylation extractor output files. Average methylation in 100 kB non overlapping windows were determined using BEDTools. Methylation correlation at individual CpG sites and at 100 kB non overlapping windows and pair-wise root mean square error (RMSE) values were computed using custom R scripts.

## Data availability

The WGBS data herein is submitted to NCBI’s short read archive (SRA) with the accession number: XXXXX

## Acknowledgments

Sequencing was performed at the SNP&SEQ Technology Platform in Uppsala, which is part of the National Genomics Infrastructure (NGI) funded by the Swedish Council for Research Infrastructures and Science for Life Laboratory. Computational analysis was performed on resources provided by the Swedish National Infrastructure for Computing (SNIC) through the Uppsala Multidisciplinary Center for Advanced Computational Science (UPPMAX).

---

**Supplementary Figure 1.**
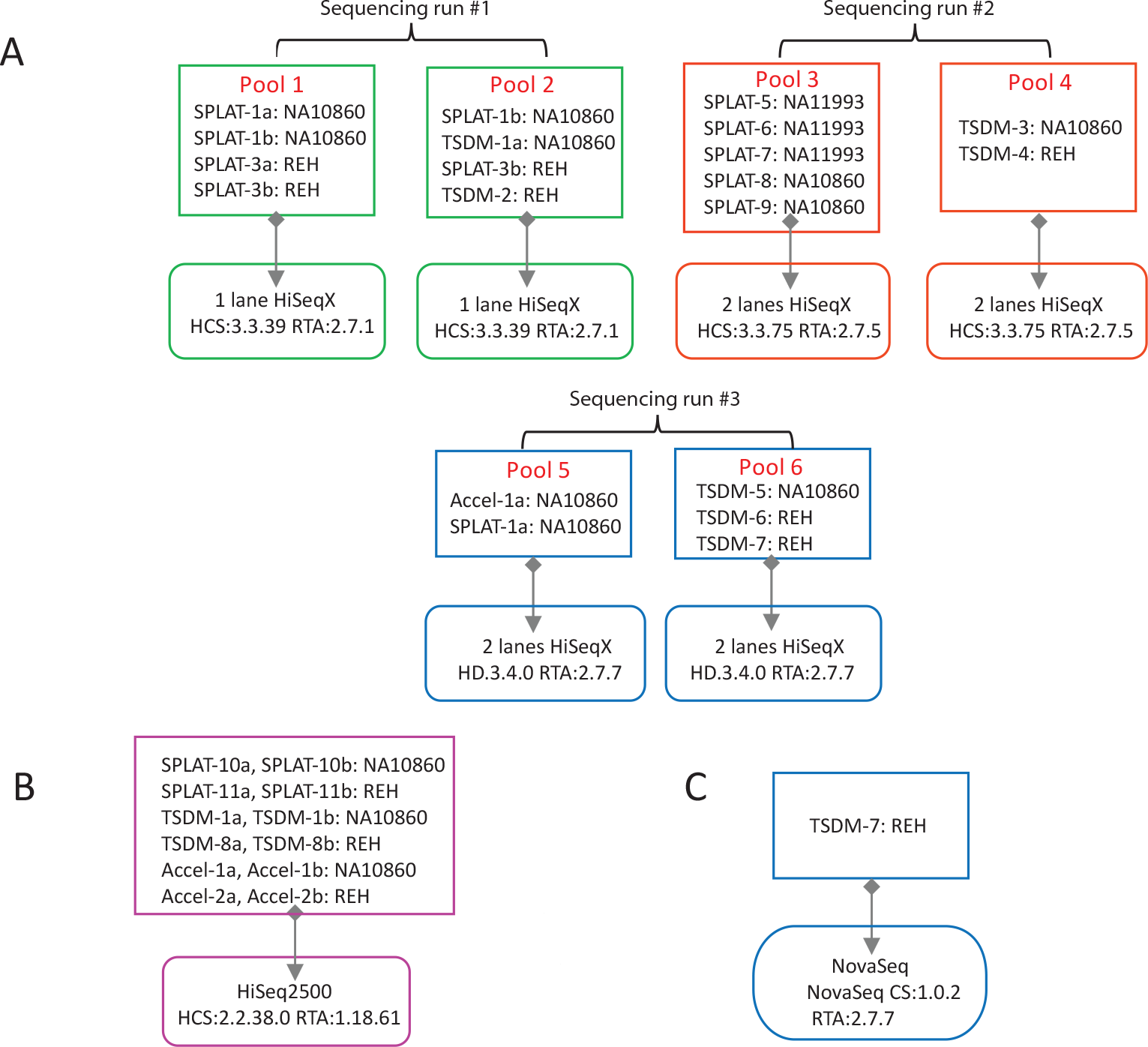
Outline of samples, library pools and software versions used in the comparison. A) Libraries sequenced on HiSeq X. For each HiSeq X sequencing run the library pooling was performed with the objective to obtain raw reads corresponding to 30x per cell line. B) Libraries sequenced on HiSeq 2500. C) Libraries sequenced on NovaSeq.

**Supplementary Figure 2.**
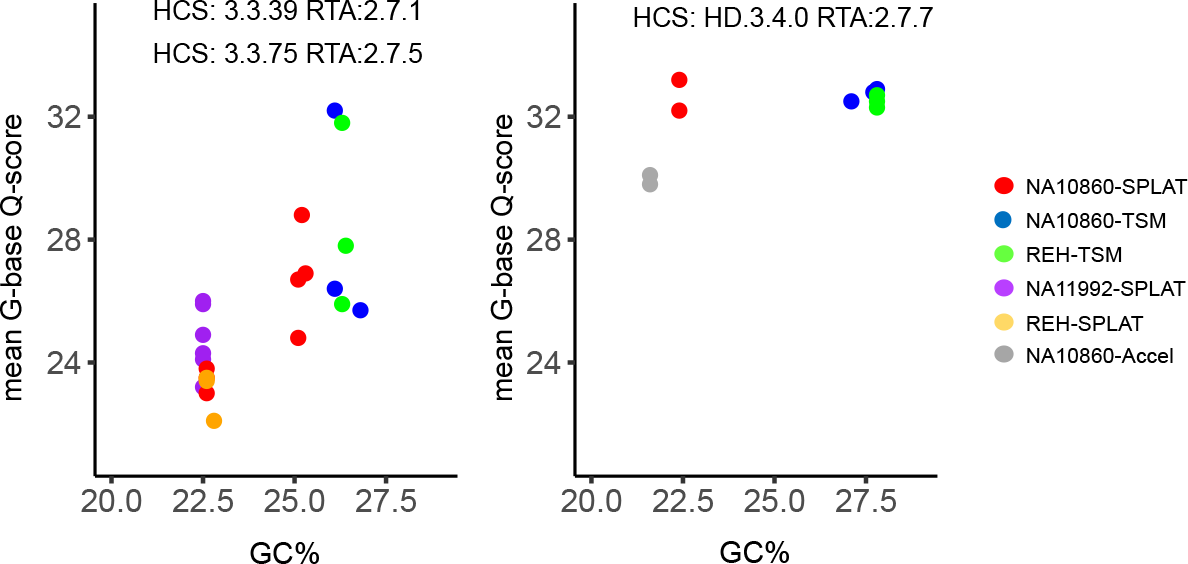
Average guanine Q-scores increases with the library GC content. For HiSeq X software versions RTA 2.7.1 and 2.7.7, a trend towards higher Q-scores with increasing GC content was observed. This trend was not apparent in data generated with RTA 2.7.7.

**Supplementary Figure 3.**
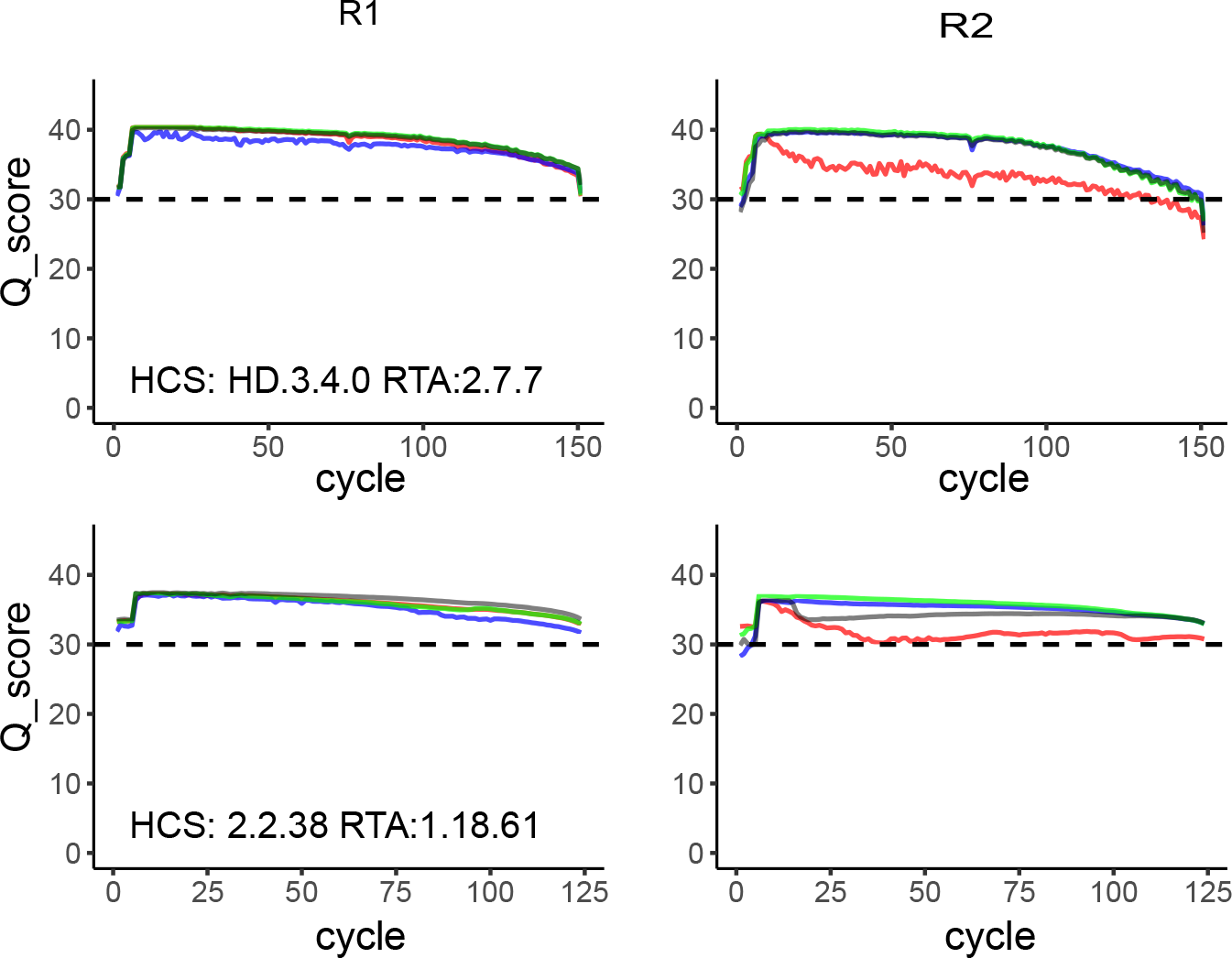
Base call quality score plots for an Accel-NGS Methyl-Seq library. Per nucleotide Q-score plots for an Accel-NGS Methyl-Seq library (cell line NA10860) sequenced both on HiSeq X (RTA 2.7.7) and HiSeq 2500 (RTA 1.18.61)

**Supplementary Table 1.**
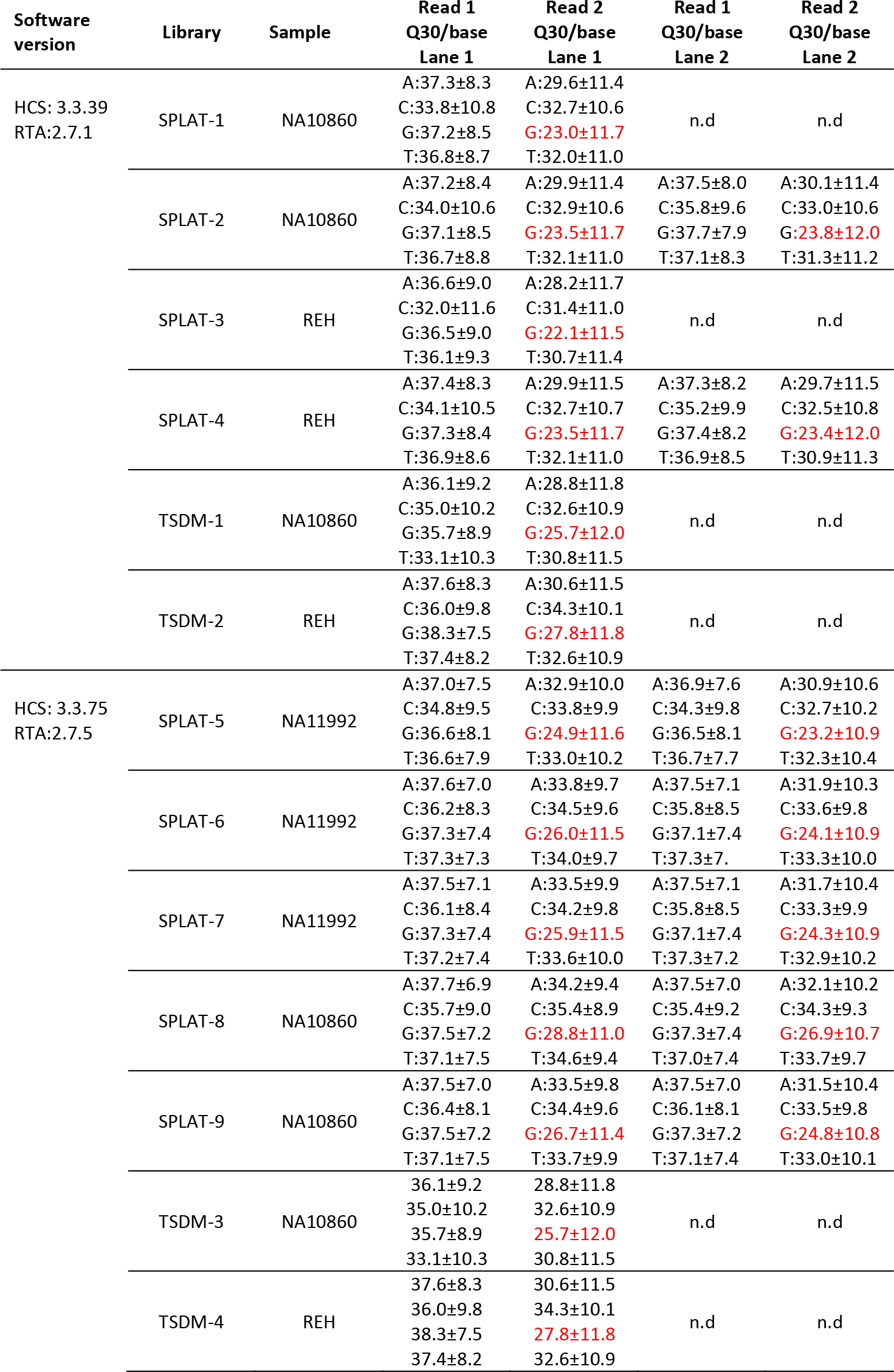
Per nucleotide quality scores for all sequencing runs performed with HCS 3.3.39/ RTA 2.7.1 and HCS 3.3.75/ RTA 2.7.5.

| Software version | Library | Sample | Read 1<br>Q30/base<br>Lane 1 | Read 2<br>Q30/base<br>Lane 1 | Read 1<br>Q30/base<br>Lane 2 | Read 2<br>Q30/base<br>Lane 2 |
| --- | --- | --- | --- | --- | --- | --- |
| HCS: 3.3.39<br>RTA:2.7.1 | SPLAT-1 | NA10860 | A:37.3±8.3 | A:29.6±11.4 | n.d | n.d |
|  |  |  | C:33.8±10.8 | C:32.7±10.6 |  |  |
|  |  |  | G:37.2±8.5 | G:23.0±11.7 |  |  |
|  |  |  | T:36.8±8.7 | T:32.0±11.0 |  |  |
|  | SPLAT-2 | NA10860 | A:37.2±8.4 | A:29.9±11.4 | A:37.5±8.0 | A:30.1±11.4 |
|  |  |  | C:34.0±10.6 | C:32.9±10.6 | C:35.8±9.6 | C:33.0±10.6 |
|  |  |  | G:37.1±8.5 | G:23.5±11.7 | G:37.7±7.9 | G:23.8±12.0 |
|  |  |  | T:36.7±8.8 | T:32.1±11.0 | T:37.1±8.3 | T:31.3±11.2 |
|  | SPLAT-3 | REH | A:36.6±9.0 | A:28.2±11.7 | n.d | n.d |
|  |  |  | C:32.0±11.6 | C:31.4±11.0 |  |  |
|  |  |  | G:36.5±9.0 | G:22.1±11.5 |  |  |
|  |  |  | T:36.1±9.3 | T:30.7±11.4 |  |  |
|  | SPLAT-4 | REH | A:37.4±8.3 | A:29.9±11.5 | A:37.3±8.2 | A:29.7±11.5 |
|  |  |  | C:34.1±10.5 | C:32.7±10.7 | C:35.2±9.9 | C:32.5±10.8 |
|  |  |  | G:37.3±8.4 | G:23.5±11.7 | G:37.4±8.2 | G:23.4±12.0 |
|  |  |  | T:36.9±8.6 | T:32.1±11.0 | T:36.9±8.5 | T:30.9±11.3 |
|  | TSDM-1 | NA10860 | A:36.1±9.2 | A:28.8±11.8 | n.d | n.d |
|  |  |  | C:35.0±10.2 | C:32.6±10.9 |  |  |
|  |  |  | G:35.7±8.9 | G:25.7±12.0 |  |  |
|  |  |  | T:33.1±10.3 | T:30.8±11.5 |  |  |
|  | TSDM-2 | REH | A:37.6±8.3 | A:30.6±11.5 | n.d | n.d |
|  |  |  | C:36.0±9.8 | C:34.3±10.1 |  |  |
|  |  |  | G:38.3±7.5 | G:27.8±11.8 |  |  |
|  |  |  | T:37.4±8.2 | T:32.6±10.9 |  |  |
| HCS: 3.3.75<br>RTA:2.7.5 | SPLAT-5 | NA11992 | A:37.0±7.5 | A:32.9±10.0 | A:36.9±7.6 | A:30.9±10.6 |
|  |  |  | C:34.8±9.5 | C:33.8±9.9 | C:34.3±9.8 | C:32.7±10.2 |
|  |  |  | G:36.6±8.1 | G:24.9±11.6 | G:36.5±8.1 | G:23.2±10.9 |
|  |  |  | T:36.6±7.9 | T:33.0±10.2 | T:36.7±7.7 | T:32.3±10.4 |
|  | SPLAT-6 | NA11992 | A:37.6±7.0 | A:33.8±9.7 | A:37.5±7.1 | A:31.9±10.3 |
|  |  |  | C:36.2±8.3 | C:34.5±9.6 | C:35.8±8.5 | C:33.6±9.8 |
|  |  |  | G:37.3±7.4 | G:26.0±11.5 | G:37.1±7.4 | G:24.1±10.9 |
|  |  |  | T:37.3±7.3 | T:34.0±9.7 | T:37.3±7.1 | T:33.3±10.0 |
|  | SPLAT-7 | NA11992 | A:37.5±7.1 | A:33.5±9.9 | A:37.5±7.1 | A:31.7±10.4 |
|  |  |  | C:36.1±8.4 | C:34.2±9.8 | C:35.8±8.5 | C:33.3±9.9 |
|  |  |  | G:37.3±7.4 | G:25.9±11.5 | G:37.1±7.4 | G:24.3±10.9 |
|  |  |  | T:37.2±7.4 | T:33.6±10.0 | T:37.3±7.2 | T:32.9±10.2 |
|  | SPLAT-8 | NA10860 | A:37.7±6.9 | A:34.2±9.4 | A:37.5±7.0 | A:32.1±10.2 |
|  |  |  | C:35.7±9.0 | C:35.4±8.9 | C:35.4±9.2 | C:34.3±9.3 |
|  |  |  | G:37.5±7.2 | G:28.8±11.0 | G:37.3±7.4 | G:26.9±10.7 |
|  |  |  | T:37.1±7.5 | T:34.6±9.4 | T:37.0±7.4 | T:33.7±9.7 |
|  | SPLAT-9 | NA10860 | A:37.5±7.0 | A:33.5±9.8 | A:37.5±7.0 | A:31.5±10.4 |
|  |  |  | C:36.4±8.1 | C:34.4±9.6 | C:36.1±8.1 | C:33.5±9.8 |
|  |  |  | G:37.5±7.2 | G:26.7±11.4 | G:37.3±7.2 | G:24.8±10.8 |
|  |  |  | T:37.1±7.5 | T:33.7±9.9 | T:37.1±7.4 | T:33.0±10.1 |
|  | TSDM-3 | NA10860 | 36.1±9.2 | 28.8±11.8 | n.d | n.d |
|  |  |  | 35.0±10.2 | 32.6±10.9 |  |  |
|  |  |  | 35.7±8.9 | 25.7±12.0 |  |  |
|  |  |  | 33.1±10.3 | 30.8±11.5 |  |  |
|  | TSDM-4 | REH | 37.6±8.3 | 30.6±11.5 | n.d | n.d |
|  |  |  | 36.0±9.8 | 34.3±10.1 |  |  |
|  |  |  | 38.3±7.5 | 27.8±11.8 |  |  |
|  |  |  | 37.4±8.2 | 32.6±10.9 |  |  |

**Supplementary Table 2.**
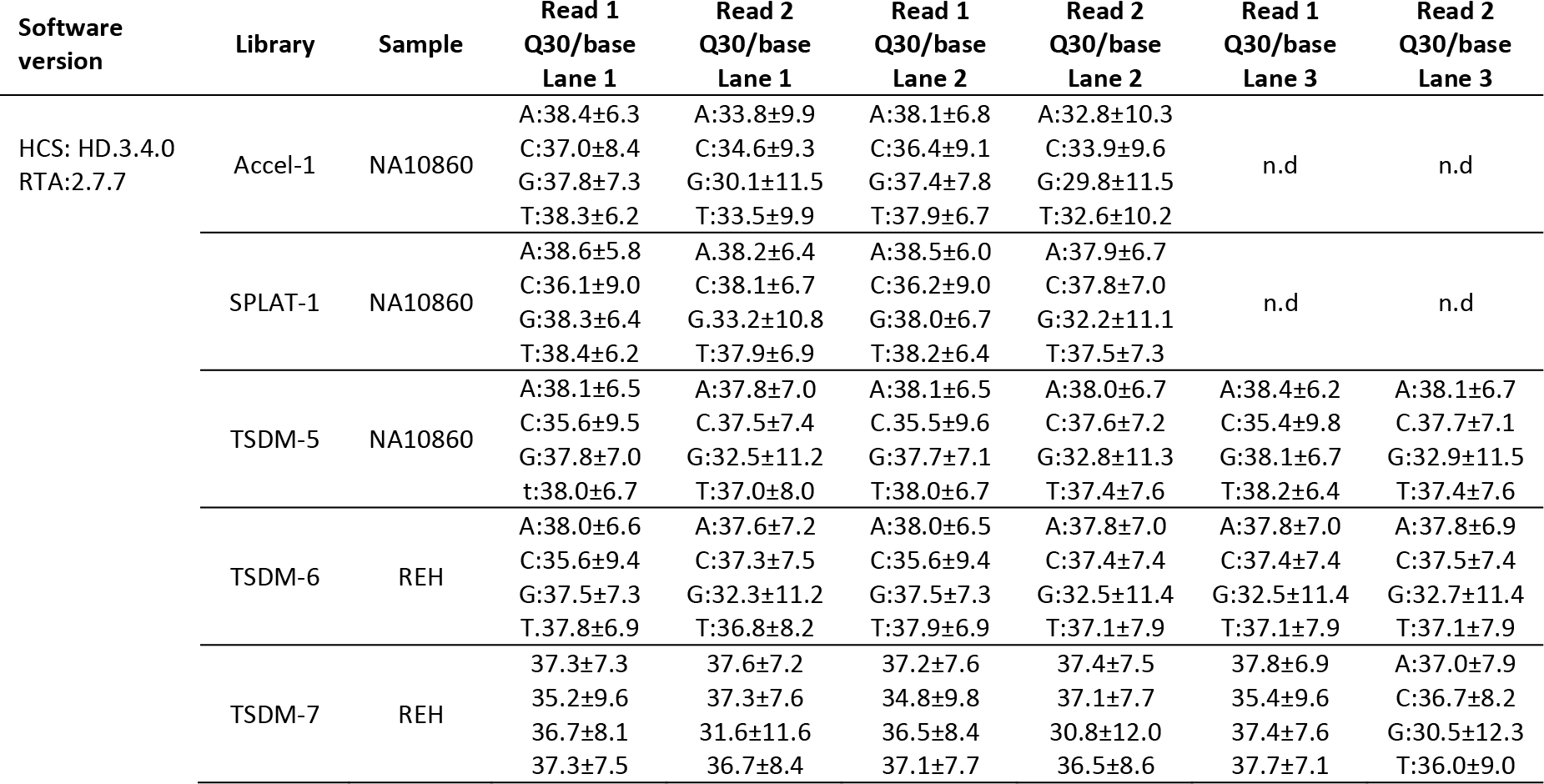
Per nucleotide quality scores for all sequencing runs performed with HCS HD 3.4.0/ RTA 2.7.7.

**Supplementary Table 3.**
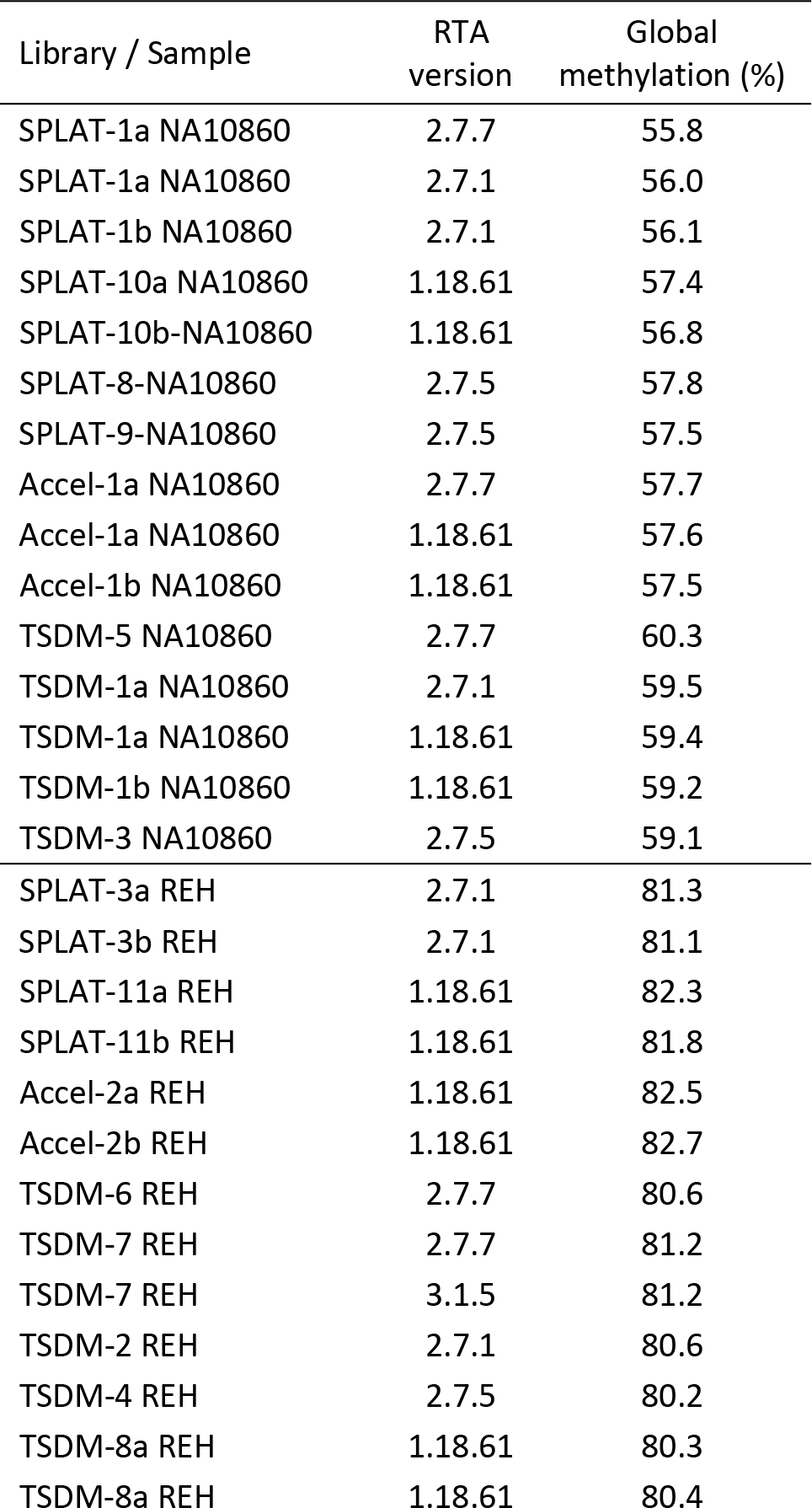
Global methylation levels per library and software version.

